# GSDMD pore formation regulates caspase-4 cleavage to limit IL-18 production in the intestinal epithelium

**DOI:** 10.1101/2024.02.01.578487

**Authors:** J.K. Bruce, L. Li, Y. Tang, N. Winsor, C. Krustev, S. Keely, D.J. Philpott, S.E. Girardin

## Abstract

Epithelial inflammasomes induce pyroptosis and release cytokines to defend against cytosolic pathogens. However, pyroptosis in epithelial barriers must be carefully regulated to facilitate elimination of infected cells while limiting widespread pyroptosis to preserve the single cell barrier. How epithelial cells achieve this is unknown. In this study, we describe a novel epithelial caspase regulation mechanism. By examining caspase-4 activation in human epithelial cells, we discovered that GSDMD pore formation serves as a signal to terminate caspase-4 activity thus facilitating epithelial cell expulsion while controlling cytokine secretion. Inhibition of epithelial pyroptosis led to IL-18 hyperproduction, likely as a mechanism to combat increased pathogen burden and initiate a wider immune response. Moreover, we demonstrate that full-length, rather than cleaved caspase-4 is active against IL-18 and propose that GSDMD pore formation facilitates cleavage of caspase-4 to terminate its catalytic activity. By comparing human cells and murine epithelial organoids to immune cells, we show that GSDMD pore mediated inhibition of caspase activity is largely specific to epithelial cells. Overall, these studies characterise a novel, epithelial-specific negative feedback loop that modulates inflammasome activity and challenge the dogma that autocatalytic caspase cleavage is required for caspase activity against substrates.

**Graphical Abstract:** In intestinal epithelial cells, caspase activation simultaneously leads to GSDMD pore formation and IL-18 release. GSDMD pore formation provides a signal to terminate caspase activity and limit cytokine production. In GSDMD deficient cells, lack of an inhibition signal leads to caspase mediated IL-18 hyperproduction. Upon cell death this leads to release of massive amounts of IL-18. Created with BioRender.com

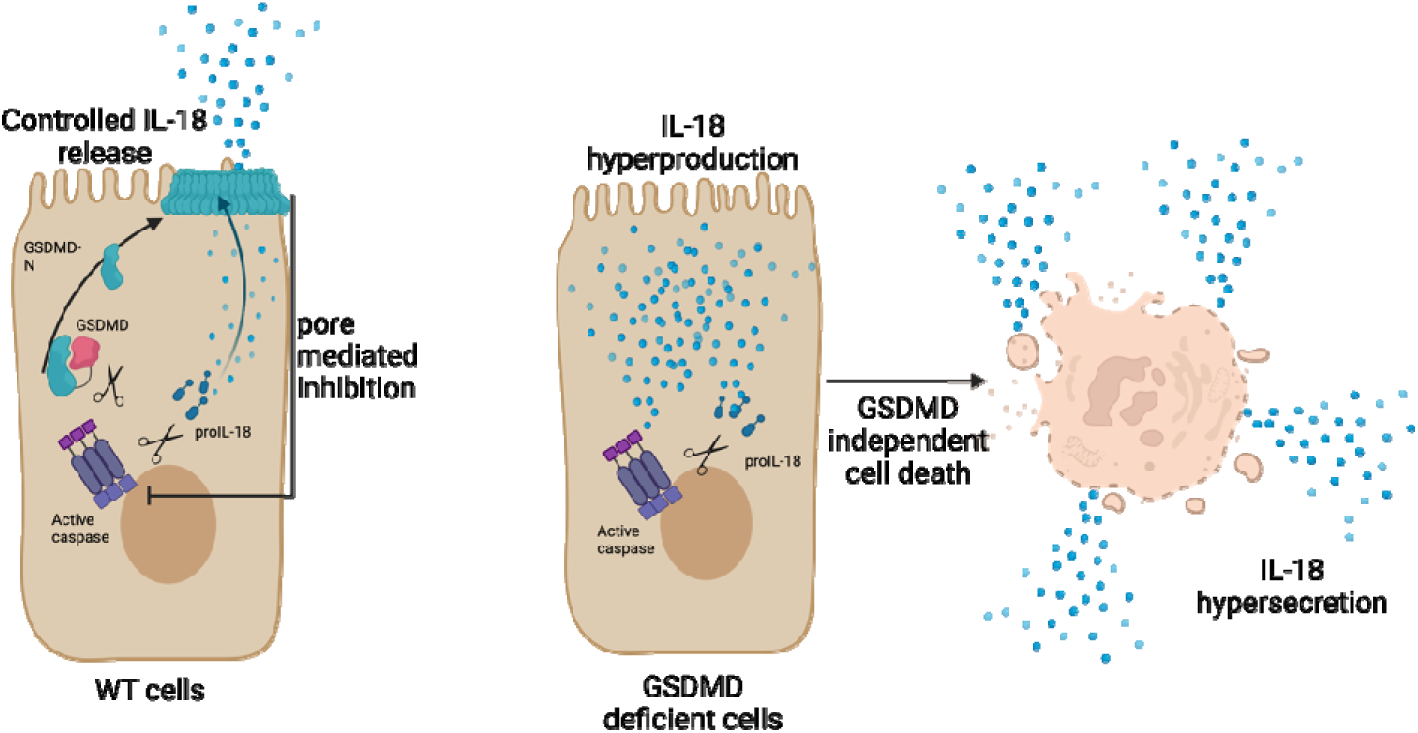

## Introduction

Inflammasomes are innate immune signalling complexes that recognise a diverse range of inflammatory molecules to activate inflammatory caspases^1^. Inflammatory caspase activation leads to cleavage of gasdermin-d (GSDMD)^2,3^ and the cytokines IL-18 and IL-1β. Cleaved GSDMD forms pores in the plasma membrane that facilitate the release of biologically active IL-18 and IL-1β^4^. GSDMD pores also lead to the inflammatory cell death known as pyroptosis^5^.

Inflammasomes are most widely studied in the context of myeloid cells. In myeloid cells, rapid lytic death of singular cells eliminates a pathogen’s replicative niche. The pyroptotic cell carcass traps bacteria and the cell releases phagocytic chemo-attractants^6^. Thus, in myeloid cells cellular lysis is an efficient mechanism to rapidly contain infections. Inflammasomes are also key defenders of epithelial barriers, where rapid lytic cell death of epithelial cells would be detrimental to host function. Pyroptosis must be tightly controlled to prevent disruptions to epithelial barrier integrity. In mice, *Salmonella* infection results in the rapid extrusion of infected epithelial cells^7,8^. During this process, barrier integrity is maintained by GSDMD- and caspase-1-dependent actin rearrangements which rapidly re-seal the epithelium around the extruded cell^8^. While caspase-1 or GSDMD deficiency results in a loss of epithelial barrier integrity, multiple redundant pathways exist to facilitate epithelial cell extrusion in the case of inhibition of NLRC4 components, including ASC-caspase-8 dependent expulsion mechanism^7^ and activation of the caspase-11 inflammasome^9^. The existence of redundant pathways indicates that co-ordinated epithelial cell pyroptosis is a critical and highly conserved pathogen defense mechanism.

In the human intestinal epithelium, expression of NLRC4 inflammasome components may not be as critical as in the murine intestinal epithelium^10^. Stimuli such as *Salmonella* and *Shigella* that in mice induce a rapid NLRC4-dependent response instead activate the intracellular LPS-sensing caspase-4 inflammasome. Caspase-4, rather than caspase-1, restricts bacterial replication and facilitates IL-18 release^10,11^. Together these data indicate that the caspase-4 inflammasome is the key responder in the human intestinal epithelium. How caspase-4 controls pyroptosis in epithelial barriers and the consequences of inhibiting epithelial cell pyroptosis in humans is unknown.

Current research efforts have placed emphasis on understanding the role of epithelial cell death and expulsion. Intestinal inflammasomes additionally activate and release IL-18. Epithelial derived IL-18 recruits neutrophils and natural killer cells to restrict intestinal pathogen dissemination^12^. Additionally, IL-18 is involved in expansion of epithelial cell stem cells^13^, and it is possible that IL-18 released during pyroptosis assists in epithelial cell regeneration to account for the expelled cells. Importantly, it has been suggested that caspase-4, rather than caspase-1 is required for IL-18 processing in the human epithelium^10,11,14^, however to our knowledge, a direct action of caspase-4 on IL-18 has not been demonstrated. Understanding how caspase-4 activates and releases IL-18 during pyroptosis is key for understanding the co-ordinated inflammasome response in the human intestinal epithelium.

Until recently it was thought that cell death terminated inflammasome signalling and cytokine production. However, studies are increasingly showing that cells have mechanisms to modulate pyroptosis and prevent cell death following inflammasome activation. GSDMD pore induced calcium influx induces the ESCRT system to remove pores from the plasma membrane to limit pyroptosis^15^. Certain cell types can secrete inflammasome dependent IL-1β for up to 72 hours while remaining viable^4,16,17^. These studies reveal that there is dynamic regulation in the induction of pyroptosis. It has been proposed that the balance of pyroptosis and cytokine release is controlled by the degree to which a cell can activate pore repair pathways^18^. However, this fails to account for a mechanism to turn-off inflammasome signalling, stop cytokine production and return to homeostasis. One study has proposed that caspase-1 cleavage at the ASC speck is a two-step process: binding to ASC induces an activating cleavage and a secondary cleavage releases caspase-1 from the ASC speck to terminate catalytic activity^19^. It is unclear what governs the rate of caspase-1 activation and inactivation. How ASC-independent non-canonical inflammasomes regulate activation is unknown. The mechanisms that fine-tune pyroptosis are unclear, however they may be particularly important at epithelial barriers where minimising cell death while facilitating cytokine release is so important for maintaining barrier integrity and orchestrating a wider immune response.

In this study, we investigated how the caspase-4 inflammasome functions in the intestinal epithelium. We demonstrate that caspase-4 directly cleaves IL-18. Moreover, we uncover that full length caspase-4 rather than cleaved caspase-4 has greater IL-18 cleaving capacity. We propose that in WT cells, GSDMD pore formation serves as a signal to cleave caspase-4 and terminate IL-18 processing hence that GSDMD pore formation constitutes an inflammasome-intrinsic feedback mechanism to terminate caspase activity. We demonstrate that this feedback mechanism has a functional significance. During infection, controlled pyroptosis and IL-18 secretion limit epithelial cell death and facilitate cytokine secretion. When GSDMD is inhibited or absent, the lack of caspase cleavage results in IL-18 hypersecretion. This may be a compensatory mechanism to promote wider immune activation to combat increased bacterial load when epithelial cell expulsion mechanisms are impaired. Overall, this work further characterises how caspase-4 functions in the intestinal epithelium and defines a new mechanism for regulation of human epithelial inflammasome signalling.

## Methods

### Cell culture

HCT116, C2bbe1 and HEK293T cells were maintained in Dulbecco’s Modified Eagle’s Medium (DMEM) (Wisent) supplemented with 10% heat-inactivated fetal bovine serum (FBS; Wisent) and 1% penicillin/streptomycin. THP1 cells were maintained in RPMI 1640 media (Wisent) supplemented with 10% heat – inactivated fetal bovine serum (FBS; Wisent) and 0.05mM 2-mercaptoethanol. Cells were maintained in 95% air, 5% CO_2_ and 37°C. Cells were used for a maximum of 10 passages.

### Generation of CRISPR knockout cells

HCT116 knockout cells were generated using the using lentiCRISPRv2 plasmid (Addgene). Guides were designed using Synthego CRISPR design tool and received from IDT. Multiple guides per target were designed. Guides were annealed then cloned into lentiCRISPRv2 plasmid and the resulting plasmid was sequenced. To generate lentivirus, lentiCRISPRv2 plasmid was transfected into HEK293T cells along with pMD2.G and psPAX2 viral envelope and packaging vectors. Virus was harvested 3 days later and used to transduce HCT116 cells. Polyclonal stably expressing cells were selected with puromycin. Monoclonal cultures were generated via the single cell limiting dilution method. Knockouts were screened by western blot and functional assays.

#### Guides used

Caspase-4 – CAC CGA GTT ATC CAA AAC ACC AGT G

GSDMD – CAC CGT CTC CGG ACT ACC CGC TCA A

Generation of shRNA stable knockdown cells

shRNA knockdowns were generated using the pLKO.1 expression vector. shRNA constructs were designed and received from IDT. shRNA constructs were annealed then cloned into pLKO.1 expression vector. The vector was sequenced. Lentivirus was generated by transfecting HEK293T cells with shRNA, pMD2.G and psPAX2 viral envelope and packaging vectors. Virus was harvested three days later. Transduced cells were selected by addition of the selection marker puromycin to culture three days after viral transduction. Knockdown of targeted protein was validated by western blot, functional assays or qPCR as indicated.

#### shRNA constructs used were

Caspase-1- CCG GCA CAC GTC TTG CTC TCA TTA TCT CGA GAT AAT GAG AGC AAG ACG TGT GTT TTT

Caspase-4 – CCG GGC AAC GTA TGG CAG GAC AAA TCT CGA GAT TTG TCC TGC CAT ACG TTG CTT TTT G

Caspase – 5 CCG GGA TTG GAT AAC TTC GTG ATA ACT CGA GTT ATC ACG AAG TTA TCC AAT CTT TTT G

GSDMD – CCG GCA ACC TGT CTA TCA AGG ACA TCT CGA GAT GTC CTT GAT AGA CAG GTT G TT TTT G

### siRNA knockdown of cells

To generate transient knockdown of targets, Silencer select siRNA (ambion life technologies) was transfected into cells using Lipofectamine RNAi MAX transfection reagent. Knockdown of target protein was validated by western blot or qPCR. Cells were used for experiments three days following transfection.

#### siRNA used

Caspase-1 – CCA CUG AAA GAG UGA CUU Utt (Thermo Fisher; siRNA ID s2408)

Caspase-4 – GAG ACU AUG UAA AGA AAG Att (Thermo Fisher; siRNA ID s2414)

GSDMD – GGA ACU CGC UAU CCC UGU Utt (Thermo Fisher; siRNA ID s36339)

### Electroporation of cells

To activate inflammasomes, cells were harvested and resuspended in 100μL of Ingenio® EZporator® Electroporation Solution (Mirus Bio). Ligand was added (2μg LPS for epithelial cells, 1μg LPS for THP1 cells) and cells were nucleofected using the Amaxa Nucleofector® 1 device. Cells were placed into fresh culture medium and left for 3 hours.

### Cell cytotoxicity assay

Cell supernatant was harvested by centrifugation at 400 x g. 25μL of supernatant was diluted in 75μL of water. Percentage cell death was measured using Cell Cytotoxicity Detection kit (LDH; Sigma) as per manufacturer’s instructions.

### Western blotting

Cells were harvested in Radioimmunoprecipitation Assay Buffer (RIPA). Protein was quantified using BCA protein assay kit (Thermo). Samples were normalised in RIPA buffer and SDS-PAGE denaturing sample buffer. SDS-PAGE was conducted using standard techniques. Gels were transferred on to polyvinylidene fluoride membrane using a semi- dry transfer apparatus and blocked using 2.5% BSA. Primary antibodies were diluted at indicated concentrations and incubated overnight at 4°C. Blots were washed 3x in 0.1% TBS-T. Secondary antibodies were diluted at a concentration of 1:10000 in 2.5% BSA in 0.1%TBS-T and incubated for 1 hour at room temperature. Blots were washed 4x then developed using SuperSignal™ West Femto Maximum Sensitivity Substrate (Thermo), or Luminata Classico Western HRP substrate (Fisher Scientific)

### ELISA

For analysis of secreted IL-18 supernatants of electroporated cells were collected and spun at 400 x g to remove cells. For measurement of total IL-18, cells and supernatant were collected and, 10μL of 33% Triton-X in PBS was added per 1mL supernatant to lyse cells into supernatant. Supernatants and lysed cell suspensions were pelleted at 10 000 x g to remove cell debris. IL-18 was measured using Human Total IL-18 DuoSet ELISA (R&D Systems) as per manufacturer’s instructions.

### qPCR

Cell pellets were washed in PBS. Total RNA was isolated using GeneJET RNA purification kit (Thermo Fisher). Genomic DNA was lysed using amplification grade DNase 1 (Invivogen) as per manufacturer’s instructions. 800ng-1μg of RNA was reverse transcribed in a mix containing M-MLV reverse transcriptase (Invivogen), dNTP mix (Biobasic), RNaseOUT™ Recombinant Ribonuclease Inhibitor (Thermo Fisher), Random Hexamer Primers (Thermo Fisher), oligoDT(18) primers (Thermo Fisher). cDNA was amplified using primers from IDT and PowerUp SYBR Green master mix (Thermo Fisher) as per manufactures instructions.

#### Primers

Caspase-1 Forward – 5’ TTT CCG CAA GGT TCG ATT TTC A

Caspase-1 Reverse – 5’ GGC ATC TGC GCT CTA CCA TC

Caspase-4 Forward – 5’ ACT TGA GGG TCT GGA CTA TAG

Caspase-4 Reverse – 5’ CCA AGA ATG TGC TGT CAG AG

GSDMD Forward – 5’ GCT GGT TAT TGA CTC TGA CTT G

GSDMD Reverse – 5’ ATC ATG GAG AGG CCA GAG

IL-18 Forward – 5’ TGC AGT CTA CAC AGC TTC GG

IL-18 Reverse – 5’ CTA CTG GTT CAG CAG CCA TCT

### FLICA assay for active caspases

FLICA assays were conducted using FAM FLICA poly caspase kit (BioRad) as per manufacturer’s instructions. Briefly, FLICA reagent was added to cells at one hour post electroporation. Cells were incubated with FLICA reagent for 30 minutes, with gentle mixing every 10 minutes. 25 minutes after the addition of FLICA, propidium iodide (PI) was added for a total incubation time of 5 minutes. Cells were washed 3x in supplied apoptosis wash buffer to wash away unbound FLICA and PI, then fixed with supplied fixative for 30 minutes. Cells were analysed on a BD LSR Fortessa (BD Bioscience) cytometer. Voltages were established using relevant controls and were maintained (+/- 10%) across replicates using calibration particles (Spherotech, URCP-38-2K). Cells were separated from debris by optical gating based off forward and side scatter area. Next, single cells were selected based off the ratio of forward scatter area to forward scatter height. Approximately 25000 single cells were captured per condition. PI^+^ cells were defined by inclusion of a positive, dead cell population, consisting of a 50:50 mixture of ethanol treated and control cells. As per established flow cytometry guidelines, all gating was based of relevant unstained (fluorescence-minus-one, FMO) populations. All data was processed in FlowJo (version 10.6).

### Plasmid mutagenesis and overexpression

pcDNA3.1 vectors expressing human caspase-4 and human IL-18 were purchased from Genscript. Mutagenesis primers were designed using Aligent Quik change primer design. Mutagenesis was conducted using QuikChangeII site directed mutagenesis kit as per manufactures instructions and mutations were validated by sequencing.

#### Mutagenesis primers

Caspase 4 C258A Forward– 5’ CATCATTGTCCAGGCCGCCAGAGGTGCAAACCGT

Caspase 4 C258A Reverse – 5’ CCCACGGTTTGCACCTCTAGCGGCCTGGACAATGATGAC

Caspase 4 D270A Forward – 5’ GAACTGTGGGTCAGAGCCTCTCCAGCATCCTTG

Caspase 4 D270A Reverse – 5’ CAAGGATGCTGGAGAGGCTCTGACCCACAGTTC

Caspase 4 D289A Forward – 5’ TCTGAGAACCTAGAGGAAGCTGCTGTTTACAAGACCCA

Caspase 4 D289A Reverse – 5’ TGGGTCTTGTAAACAGCAGCTTCCTCTAGGTTCTCAGA

### Transient transfections

1μg of each plasmid or pcDNA3.1 empty vector was reverse transfected into 1x10^6^ HEK293T cells using FUGENE HD (Promega) as per manufactures instructions. Cells were harvested 16 hours post transfection.

### Organoid culture

Colons were dissected from WT and GSDMD^-/-^ littermates, washed 3x in PBS, then incubated in PBS containing 2mM EDTA for 30 minutes with gentle agitation. Colons were placed in 50mL tubes containing fresh PBS then shaken vigorously to dislodge crypt units. Cell/crypt suspensions were strained through a 70μM filter. Crypts were washed three times in PBS containing 10% FBS. Then plated into Matrigel domes (Cultrex PathClear reduced growth factor BME, type 2; R&D Systems) in a 24 well plate. Cultures were maintained in media containing 50% WRN conditioned media supplemented with 1mg/mL primocin (InvivoGen), 1mM N-Acetylcysteine, 1x B27 supplement (Gibco) and 1x N2 supplement (Gibco) and 0.1μg/mL EGF (Thermo Fisher). Cultures were passaged every 6 days and kept in a cycle of 3 days in 5μM ROCK inhibitor (Y-27632 dihydrochloride; Tocris), 3 days in the absence of rock inhibitor, to promote growth and differentiation.

### *Salmonella* infections

*Salmonella typhimurium* was sub-cultured in Luria Burtani (LB) broth from overnight cultures until culture reached OD=600. Salmonella cultures were washed 1x in PBS then added to cells at MOI=50. Cells were centrifuged for 5min at 1500rpm then returned to the incubator for 1 hour. Following this, cells were washed 3x in PBS then media containing gentamycin (Gibco) at a final concentration of 50μg/mL was added to kill extracellular bacteria. Cells were harvested 20 hours post infection.

### Organoid Inflammasome Stimulation

On day 6 following passaging, organoids were removed from Matrigel and broken into smaller cell clusters with TrypLE Express (Gibco). Cell clumps were electroporated using Mouse/Rat Hepatocyte NucleofectorTM Kit (Lonza) and Amaxa Nucleofector® 1 device in the presence or absence of flagellin (1μg) or LPS (5μg). Cells were plated in 200μL of fresh media for 2 hours.

### Antibodies and other reagents

Rabbit anti-GSDMD (HPA044487; Sigma), mouse anti - Caspase-4 (M029-3, Marine BL), rabbit anti-caspase-1 (3866S; Cell Signalling Technology), rabbit anti-Caspase-5 (ab40887; abcam), goat anti-IL-18 (AF2548; Novus Biologicals), LPS (14011S; CST), human IFNγ (80385S; Cell Signalling Technology), Nigericin (N7143; Sigma), Flagellin (tlrl-epstfla; InvivoGen), Z-VAD-FMK (tlrl-vad; InvivoGen), VX-765 (7143/50, Cedarlane), peroxidase conjugated anti-rabbit and anti-mouse secondaries (Jackson Labs, 111-035-003), peroxidase conjugated anti-goat secondary (HAF109; R&D Systems).

### Statistics

Prism software was used to plot data and determine statistical significance using analysis of variance (ANOVA) with multiple comparisons Data are presented as S.E.M as indicated. A p value of 0.05 or less was considered statistically significant.

## Results

### Loss of GSDMD results in IL-18 hyperproduction in human epithelial cells

To activate the non-canonical inflammasome, we electroporated LPS into human epithelial HCT116 cells. LPS electroporation led to cell death (Figure 1a), GSDMD cleavage (Figure 1b) and IL-18 release (Figure 1c) that was dependent on caspase-4 and GSDMD expression. Unexpectedly, when examining IL-18 production by western blot, we observed high levels of mature IL-18 in the lysates of GSDMD^-/-^ cells (Figure 1d). Mature IL-18 is secreted through GSDMD pores; therefore, to compare the total amount of IL-18 produced we lysed cells into the supernatant and measured IL-18 (hereafter referred to as total IL-18) via ELISA. Surprisingly, we observed that GSDMD^-/-^ cells produced approximately 10 times more mature IL-18 than WT cells (Figure 1e). This increase in IL-18 production could not be accounted for by elevated expression of the pro-IL-18, (Figure 1f), nor could it be inhibited by blocking transcription (Figure 1g) or translation of the pro IL-18 substrate (Figure 1h). Thus, human HCT116 intestinal epithelial cells lacking GSDMD produce increased amounts of mature IL-18 in response to caspase-4-dependent detection of intracellular LPS.

**Figure 1:**
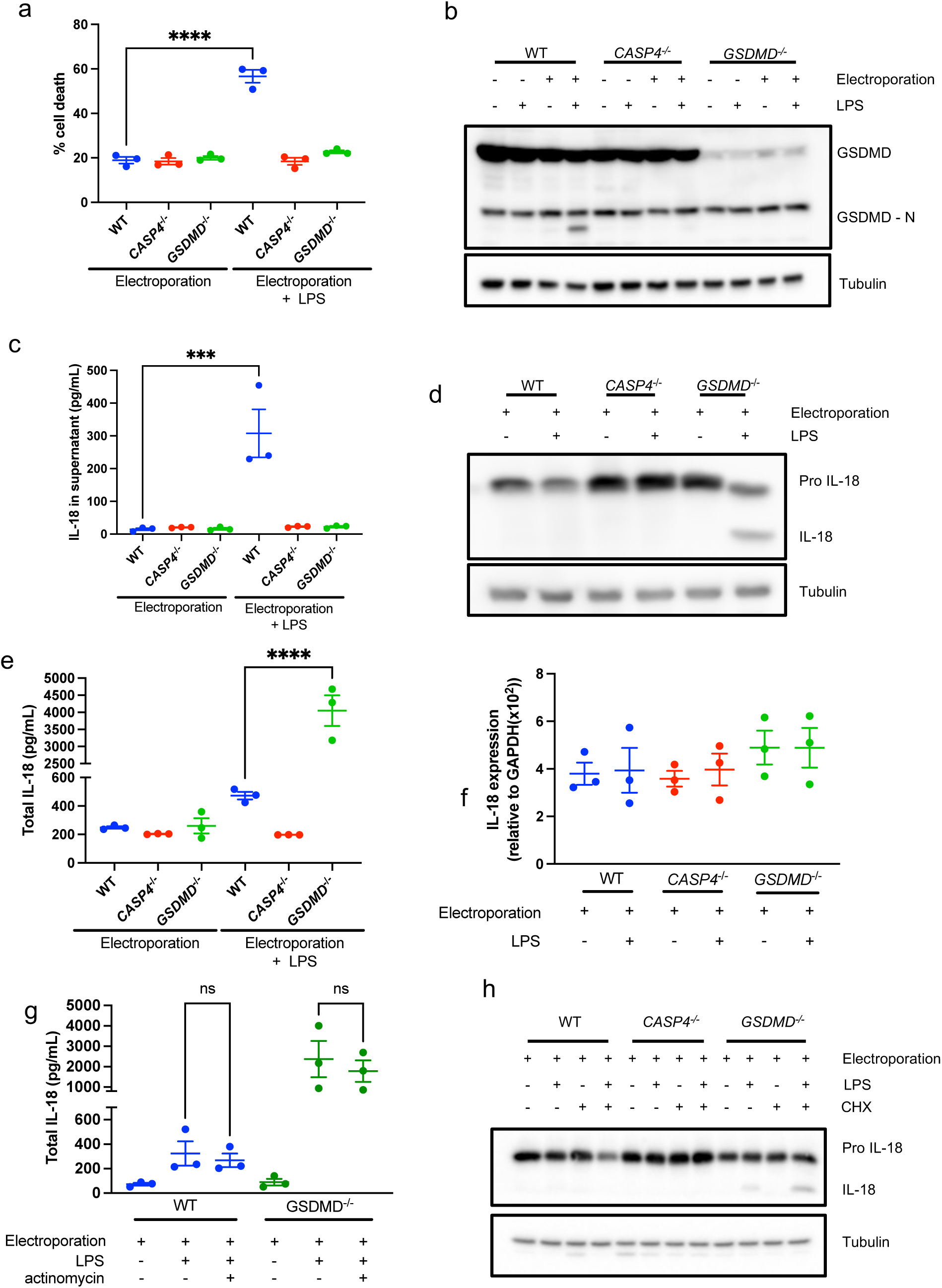
GSDMD limits production of mature IL-18 in human intestinal epithelial cells. HCT116 human epithelial cells were electroporated with 2µg LPS as indicated. Cells were harvested 3 hours post electroporation. Cell death (a) was measured using LDH cytotoxicity assay. GSDMD (b) and IL-18 cleavage (d) were measured by western blot of whole cell lysates. IL-18 secretion was measured in cell free supernatant (c). Total IL-18 (e, g) was measured by lysing cells into supernatant then conducting ELISA. IL-18 transcript levels were measured by qPCR (f). To block transcription, cells were pre-treated with actinomycin for 1 hour prior to electroporation (g). To block translation, cells were incubated with cycloheximide (CHX) for 6 hours prior to electroporation. Representative of at least 3 experiments, data displayed as mean ± S.E.M. ***p<0.005, ***p<0.001.

To confirm that our phenotype was not due to off target effects following CRISPR cas9 genome editing, we confirmed these findings in a Caco2 C2bbe1 cells using RNAi techniques (Figure 2a-d). To see if our findings occurred in other cell types, we knocked GSDMD down in human THP1 monocytes (Figure 2e). Knocking down GSDMD significantly reduced LPS-electroporation induced LDH release (Figure 2f) and slightly increased IL-18 production (Figure 2g-h), however the degree to which silencing GSDMD affected IL-18 production in monocytes was minimal compared to that observed in epithelial cells (Figure 2d compared to Figure 2g). Based on these results, we concluded that GSDMD regulates the production of IL-18 following non-canonical inflammasome activation via an unknown mechanism.

**Figure 2:**
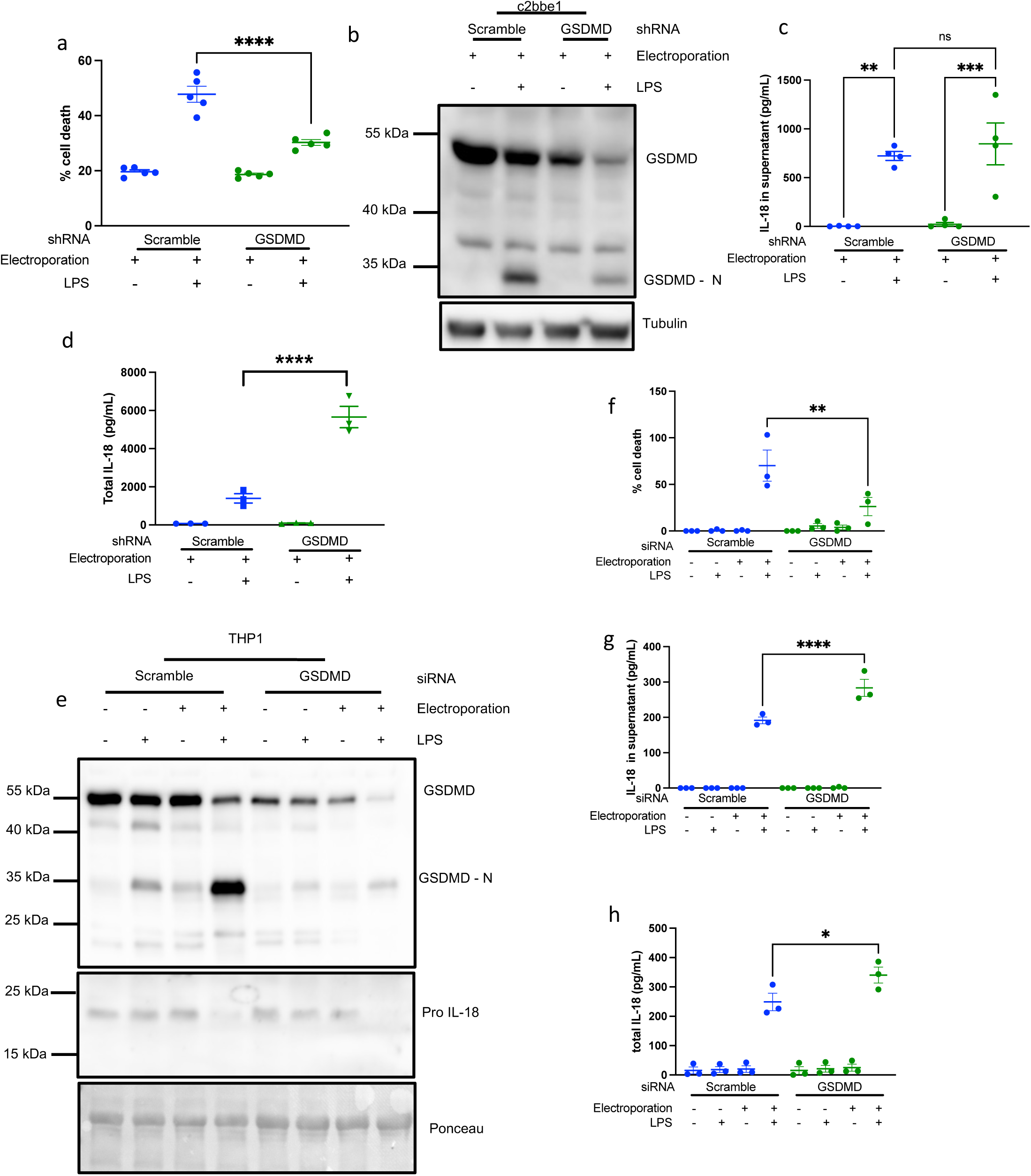
Overproduction of IL-18 is specific to epithelial cells. C2bbe1 human epithelial cells (a-d) and THP1 human monocytes (e-f) were electroporated with 2µg LPS as indicated. Cell death was measured by LDH assay (a, f), GSDMD expression and cleavage was measured by western blot (b, e), IL-18 in the supernatant was measured by ELISA (c, g), total IL-18 (d, h) was measured by lysing cells into supernatant prior to ELISA. Representative of at least 3 experiments, data displayed as mean ± S.E.M. ****p<0.001, **p<0.01, *p<0.05.

### Epithelial cell IL-18 hyperproduction occurs in the absence of LPS-induced caspase cleavage

It is widely reported that oligomerization-induced autoproteolysis is required for caspase activity against substrates^19^. Indeed, measuring the production of caspase-1 p20 is the gold standard for assessing caspase-1 activation, and increased cleavage is thought to reflect a greater degree of caspase activation. Caspase-4 is reported to be cleaved into p31 and p33 fragments^20^, and caspase-4, rather than caspase-1 is required for IL-18 production in human intestinal epithelial cells. (Supplementary Figure 1, refs^10,11,14^). We therefore hypothesized that increased caspase-4 cleavage led to increased IL-18 production in GSDMD deficient cells. LPS electroporation induced a reduction of pro-caspase-4 levels in the lysate of WT epithelial cells, and the appearance of two cleavage fragments in the supernatant (Figure 3a, b), however no LPS inducible cleavage fragment was visible in *GSDMD*^-/-^ or GSDMD KD epithelial cells (Figure 3a, b) and levels of pro-caspase-4 remained consistent following LPS electroporation. We observed a caspase-4 p31 fragment was enriched in lysates of both WT and GSDMD knockdown THP1 cells following LPS electroporation (Figure 3c). The appearance of caspase-4 p31 correlated with similar levels of IL-18 production in WT and GSDMD KD THP1 cells (as previously shown in Figure 2h). Interestingly, generation of caspase-4 p31 was blocked by pre-treatment with the NLRP3 inhibitor MCC950 (Figure 3d), indicating that this cleavage fragment is likely the result of NLRP3 activation rather than caspase-4 autoproteolytic activity. Given that there was no LPS-inducible difference in caspase-4 processing in GSDMD deficient epithelial cells we considered that loss of GSDMD might induce caspase-4-independent IL-18 production. Caspase-1 expression is low in human intestinal epithelial cells (Supplementary Figure 1f), and while LPS electroporation induced robust caspase-1 processing in THP1 monocytes, no caspase-1 activation was observed in epithelial cells (Figure 3e). LPS electroporation induced caspase-5 processing in WT HCT116 epithelial cells, however this cleavage was absent in *GSDMD^-^*^/-^ cells. Thus, in epithelial cells, a lack of inflammatory caspase cleavage is associated with IL-18 hyperproduction.

**Figure 3.**
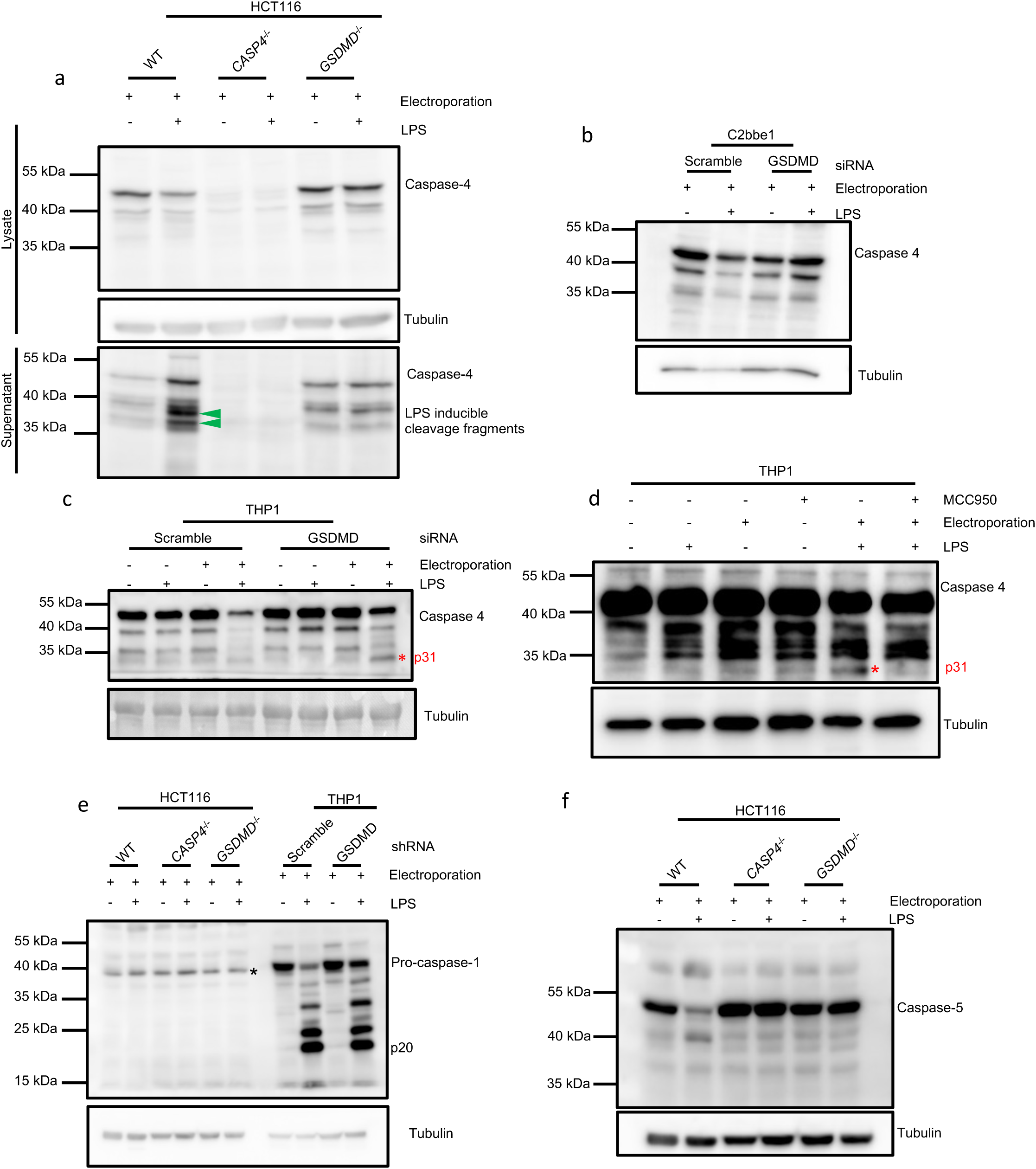
Epithelial cell IL-18 hyperproduction occurs without LPS inducible inflammatory caspase cleavage, compared to human monocytes. Human epithelial cells (HCT116 and C2bbe1) and human THP1 monocytes were electroporated with 2µg LPS. For (d) THP1 cells were co-treated with NLRP3 inhibitor MCC950. Inflammatory caspase activation was measured by western blot on whole cells lysates and supernatant. For (e) star indicates non-specific band. See supplemental Figure 2. Representative of at least 3 experiments.

### Caspase-4 cleavage limits IL-18 production in intestinal epithelial cells

Given that we did not observe inflammatory caspase processing in *GSDMD*^-/-^ cells, we wondered whether IL-18 processing was dependent on caspase activity, or if an alternative pathway was engaged. To test this hypothesis, we pre-treated cells with the pan-caspase inhibitor Z-VAD-FMK for 1 hour prior to electroporation to block the activity of all caspases. Z-VAD-FMK treatment completely inhibited LPS induced cell death (Figure 3a), IL-18 release (Figure 3b) and blocked IL-18 maturation in both WT cells and *GSDMD*^-/-^ cells (Figure 3c), confirming that caspase catalytic activity is required for IL-18 processing in *GSDMD*^-/-^ cells even in the absence of LPS inducible caspase-1, 4 or 5 cleavage.

To further elucidate which caspase processes IL-18 in *GSDMD*^-/-^ cells we used RNAi techniques to knock down inflammatory caspases in GSDMD^-^deficient cells (Figure 4d, f, and h). Knocking down caspase-4 (Figure 4e) but not caspase-1 (Figure 4g) or caspase-5 (Figure 4i), blocked IL-18 production in both WT and GSDMD^-^deficient cells, confirming that caspase-4 and no other inflammatory caspases were required for IL-18 processing. However, it is important to note that caspase-4 functions as both the sensing protein and the executioner protein, and as such caspase-4 KD cells are unable to respond to intracellular LPS. To address this issue, we pre-treated cells with the caspase-1/4 catalytic inhibitor VX-765 (Figure 4j-l). In this model, cells can bind LPS via the unaffected CARD domain, but caspase-4 is rendered catalytically dead by modification of the catalytic cystine^21^. Pre-treatment with VX-765 completely blocked GSDMD cleavage (Figure 4j), IL-18 secretion (Figure 4k) and IL-18 maturation (Figure 4l), confirming that caspase-4 catalytic activity is required for IL-18 processing in both WT and GSDMD-deficient human intestinal epithelial cells and caspase-4 cleavage is associated with reduced IL-18 production.

**Figure 4:**
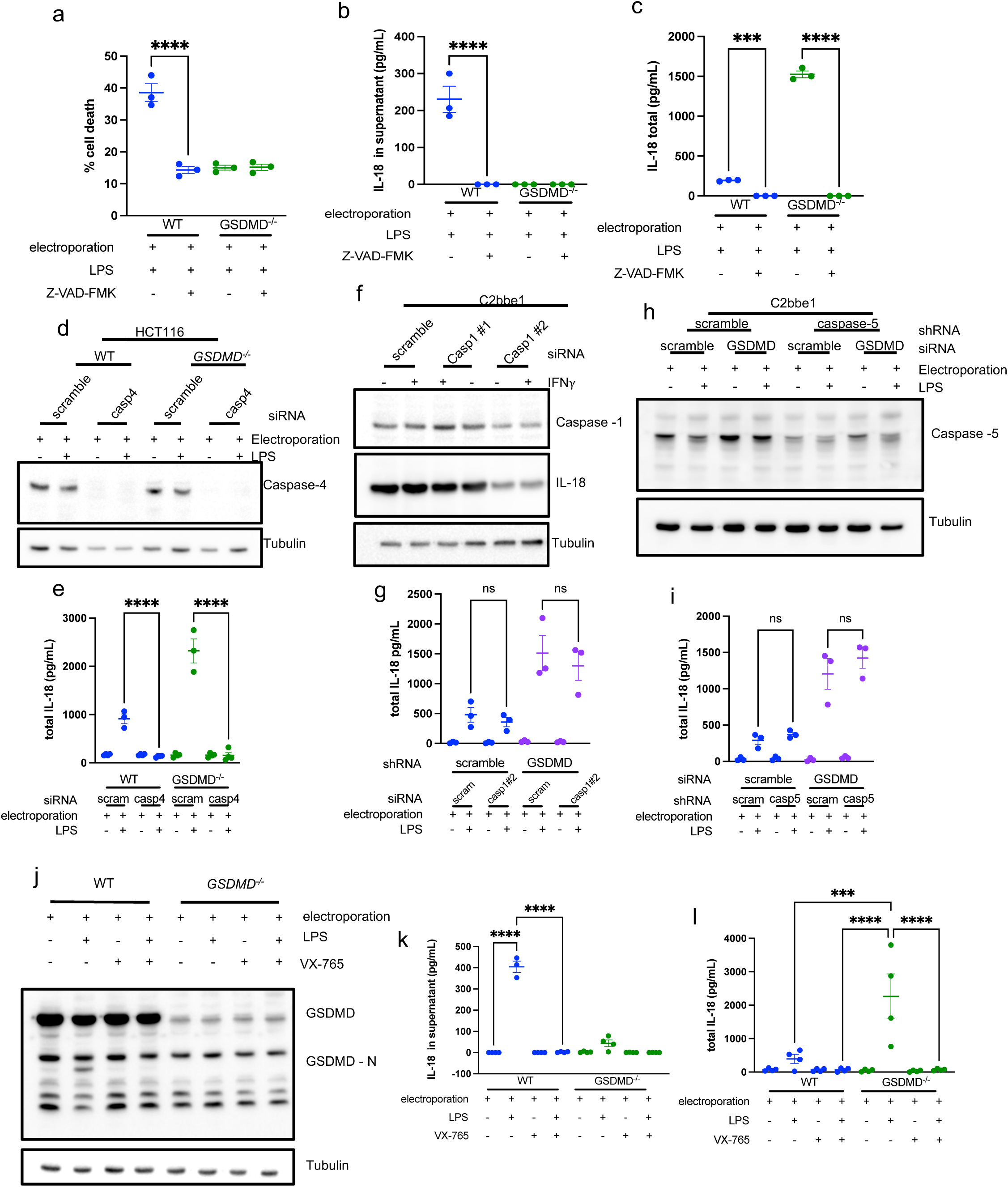
Human intestinal epithelial IL-18 production requires caspase-4 catalytic activity, but not caspase-4 cleavage. HCT116 epithelial cells were treated with 2µM of pan-caspase inhibitor Z-VAD-FMK (a-c), siRNA targeting caspase-4 (d,e) or with caspase-1/4 specific inhibitor VX-765 (j-l). C2bbe1 epithelial cell double caspase-1 or 5 and GSDMD knockdowns were generated as indicated (f-i). For inflammasome activation, cells were electroporated with 2µg LPS. Cell death was measured by LDH cytotoxicity assay(a). Western blots to assess inflammasome activation were conducted on whole protein lysates (d, f, h, j). IL-18 release was measured by ELISA on supernatant alone (b, k). Total IL-18 production was measured by lysing cells into supernatant prior to ELISA (c, e, g, i, l). Representative of at least 3 experiments, data displayed as mean ± S.E.M. ****p<0.001, ***p<0.005.

Since our data indicated that full length caspase-4 produces more IL-18, we reasoned that full length caspase-4 was likely the catalytically active species and GSDMD-facilitated cleavage might terminate activity against IL-18. We attempted to measure caspase activity via the Fluorescent Linked Inhibitor of Caspase Activity (FLICA) assay. As expected, we observed an approximately 30% increase in FLICA retention in WT cells following LPS electroporation (Figure 5a, b). LPS-induced FLICA retention was undetectable in *CASP4*^-/-^ cells (Figure 5a, b). Surprisingly, we observed no increase in FLICA retention in *GSDMD*^-/-^ cells (Figure 5a, b). We suspect that the lack of FLICA binding in *GSDMD*^-/-^ cells likely reflects that caspase-4 is in an altered, uncleaved confirmation and this increases activity against IL-18. To further investigate which caspase-4 species processes IL-18 we generated caspase-4 cleavage mutant expression vectors and transiently overexpressed these together with IL-18 and in HEK293T cells (Figure 5c, d), which do not express endogenous inflammasome components. In this system, overexpression facilitates caspase oligomerization and activation. As expected, caspase-4 was required for IL-18 processing, and mutation of the catalytic cysteine C258 prevented IL-18 cleavage. Caspase-4 is reported to auto-cleave at D270 and D289A^20^. Mutation of individual autocleavage sites slightly increased IL-18 processing compared to WT, however mutation of both sites greatly increased IL-18 processing (Figure 5d), demonstrating that non-cleavable caspase-4 produces more IL-18. We therefore propose that oligomerization of caspase-4 is sufficient to generate IL-18 processing capacity and that cleavage of the caspase limits IL-18 processing. In an endogenous system, GSDMD provides a trigger to cleave caspase-4 and terminate inflammasome activity.

**Figure 5:**
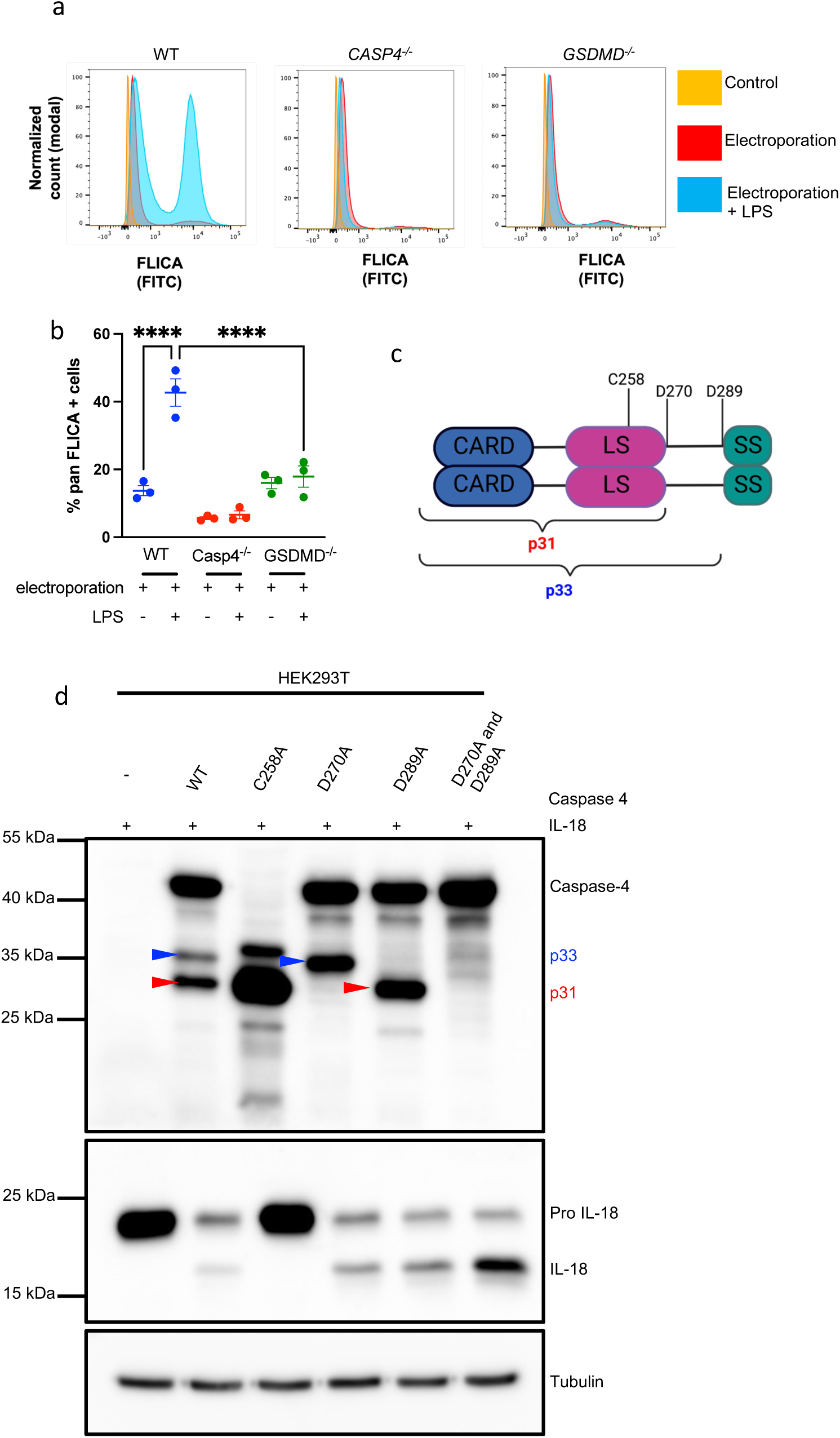
Caspase-4 cleavage limits IL-18 production. Protein in the supernatant of HCT116 cells following electroporation was extracted by TCA precipitation, caspase-4 levels were analyzed by western blot. Green arrow indicates LPS induced caspase-4 cleavage product, calculated to be 38.3 kDa (a). FLICA retention in HCT116 cells following electroporation was measured by flow cytometry (c-d). Cells were counterstained with Propidium iodide (PI) (b). Schematic of key catalytic site (C258) and known cleavage sites (D270 and D289) in caspase-4 (e). WT and indicated caspase-4 mutants were transiently overexpressed in HEK293T cells. Caspase-4 and IL-18 cleavage was measured in western blots of whole cell lysates (f). Representative of at least 3 experiments, data displayed as mean ± S.E.M. ****p<0.001, *p<0.05.

### GSDMD pore formation facilitates caspase-4 cleavage to limit IL-18 processing

We next sought to determine which function of GSDMD regulates IL-18 production and caspase-4 cleavage. We reasoned that cell lysis, GSDMD pore formation, or an effect of GSDMD cleavage on caspase-4 could each impact IL-18 processing. We first tested the effect of inhibiting pyroptotic lysis by pre-treating cells with glycine^22^. Glycine reduced cell lysis (Figure 6a) but had no effect on GSDMD cleavage (Figure 6b). Secreted IL-18 was equivalent between glycine and non-glycine treated cells, confirming that pore formation was intact (Figure 6c). Importantly, glycine had no effect on total IL-18 production (Figure 6d). Interestingly, in glycine treated cells, cleaved caspase-4 was detected in cell lysates, demonstrating that secretion of cleaved caspase-4 in WT cells does not limit IL-18 production. Together these findings demonstrate that pyroptosis does not regulate caspase-4 cleavage and IL-18 production. We next sought to determine if GSDMD pore formation regulates IL-18 processing and caspase-4 cleavage. To assess this, we pretreated cells with dimethylfumerate (DMF). DMF has been reported to succinate cysteine residues to prevent GSDMD N-terminal insertion into the plasma membrane^23^. DMF completely inhibited LPS-dependent cell lysis (Figure 7a) but had no effect on GSDMD cleavage (Figure 7b). DMF also prevented IL-18 release into the supernatant (Figure 7c). Together these results indicate that that DMF primarily inhibits GSDMD pore formation, rather that caspase-4 mediated GSDMD cleavage. Importantly, blocking GSDMD pore formation prevented caspase-4 cleavage and increased total IL-18 levels. These findings demonstrate that GSDMD pore formation provides a signal to cleave caspase-4 terminate IL-18 processing.

**Figure 6:**
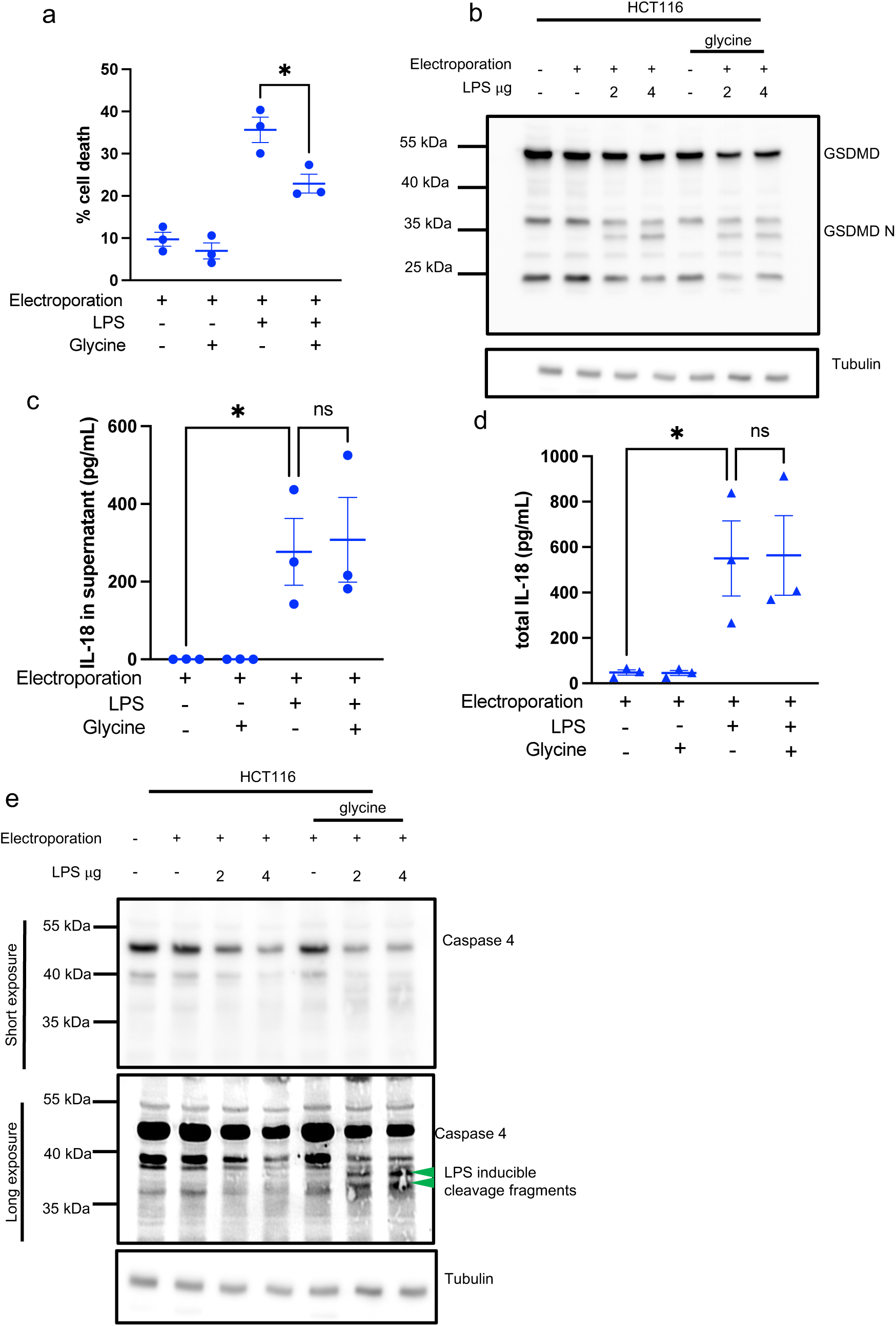
Cell lysis does not regulate caspase-4 cleavage and IL-18 production. HCT116 cells were electroporated then placed in media with 10mM glycine to prevent cell lysis (a-e). Cell death was measured by LDH cytotoxicity assay (a), GSDMD (b) and caspase-4 (e) cleavage was measured by western blots of whole cell lysates. IL-18 was measured by ELISA on supernatants (c), or on cells lysed into supernatant (d). Green arrow indicates LPS dependent caspase-4 cleavage products. Representative of at least 3 experiments, data displayed as mean ± S.E.M. *p<0.05.

**Figure 7:**
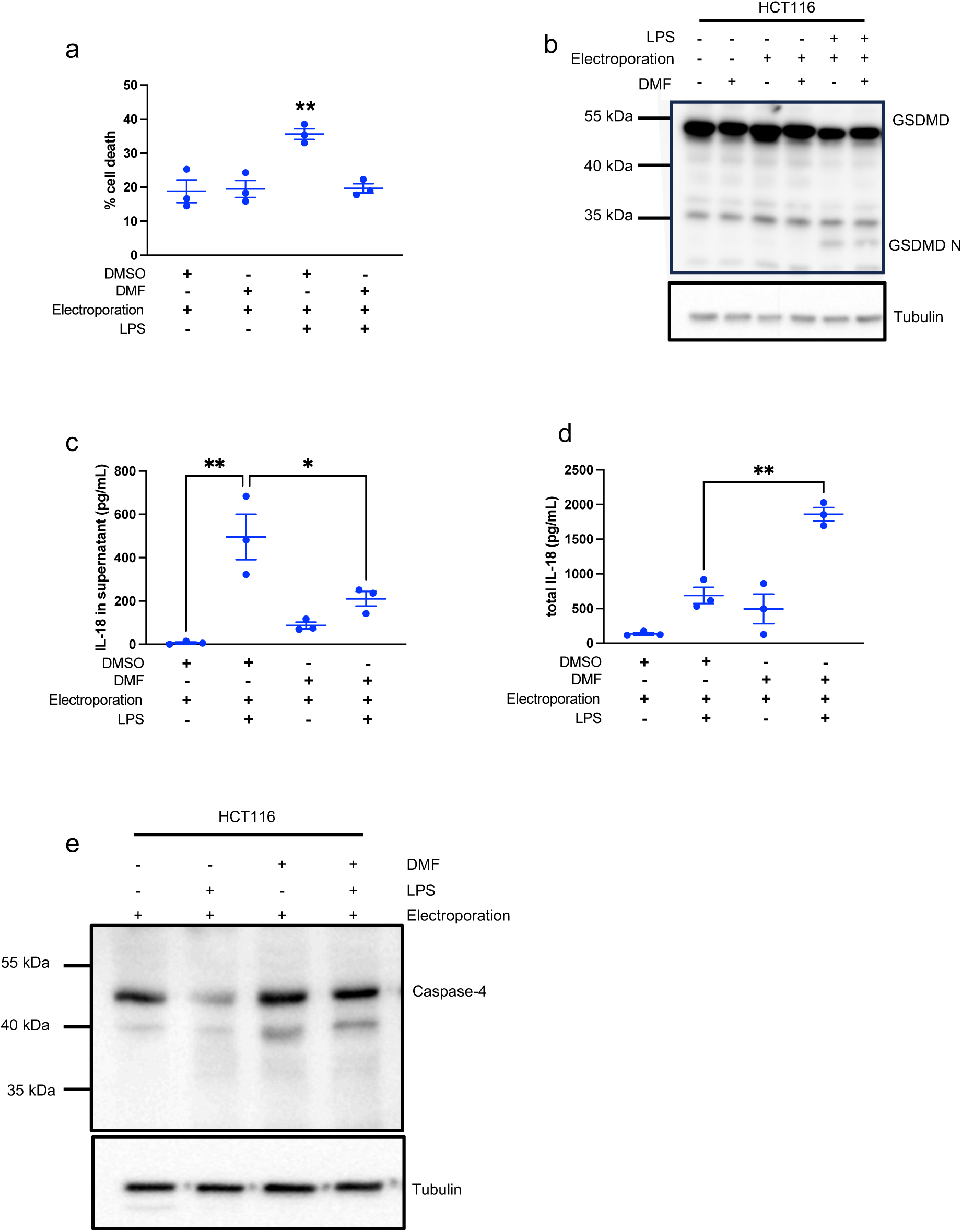
Blocking GSDMD pore formation inhibits caspase-4 cleavage and increases IL-18 production. HCT116 cells were pre-treated with 50µM dimethylfumerate (DMF) for 1 hour then electroporated with LPS and placed back into media containing 50µM DMF (a-e). Cell death was measured by LDH (a), GSDMD, and caspase-4 cleavage was measured by western blots on whole cell lysates (b, e). IL-18 was measured by ELISA on supernatant alone (c), or on cells lysed into supernatant (d). Representative of at least 3 experiments, data displayed as mean ± S.E.M. **p<0.01, *p<0.05.

### GSDMD regulates IL-18 production in murine epithelial cells following NLRC4 activation

We next wondered whether GSDMD might regulate caspase activity in other cell types and in the context of other inflammasome stimuli. To investigate whether GSDMD regulates caspase activity in murine epithelium we generated epithelial organoids and electroporated these with inflammasome stimulating ligands. Previous work has demonstrated that caspase-11 (the murine ortholog of caspase-4) does not directly process IL-18^14^. *Nlrp3* is not expressed in the murine intestinal epithelium and therefore activation of the non-canonical inflammasome in the murine epithelium stimulates GSDMD cleavage but not cytokine processing (Figure 8a). Flagellin robustly activates NLRC4 in the murine epithelium^7,24^. As with human epithelial cells, we observed increased levels of mature IL-18 in *Gsdmd*^-/-^ organoids (Figure 8b). Overall, our data demonstrates that GSDMD regulates the activity of inflammatory caspases in the epithelium in the context of different species and different caspases.

**Figure 8:**
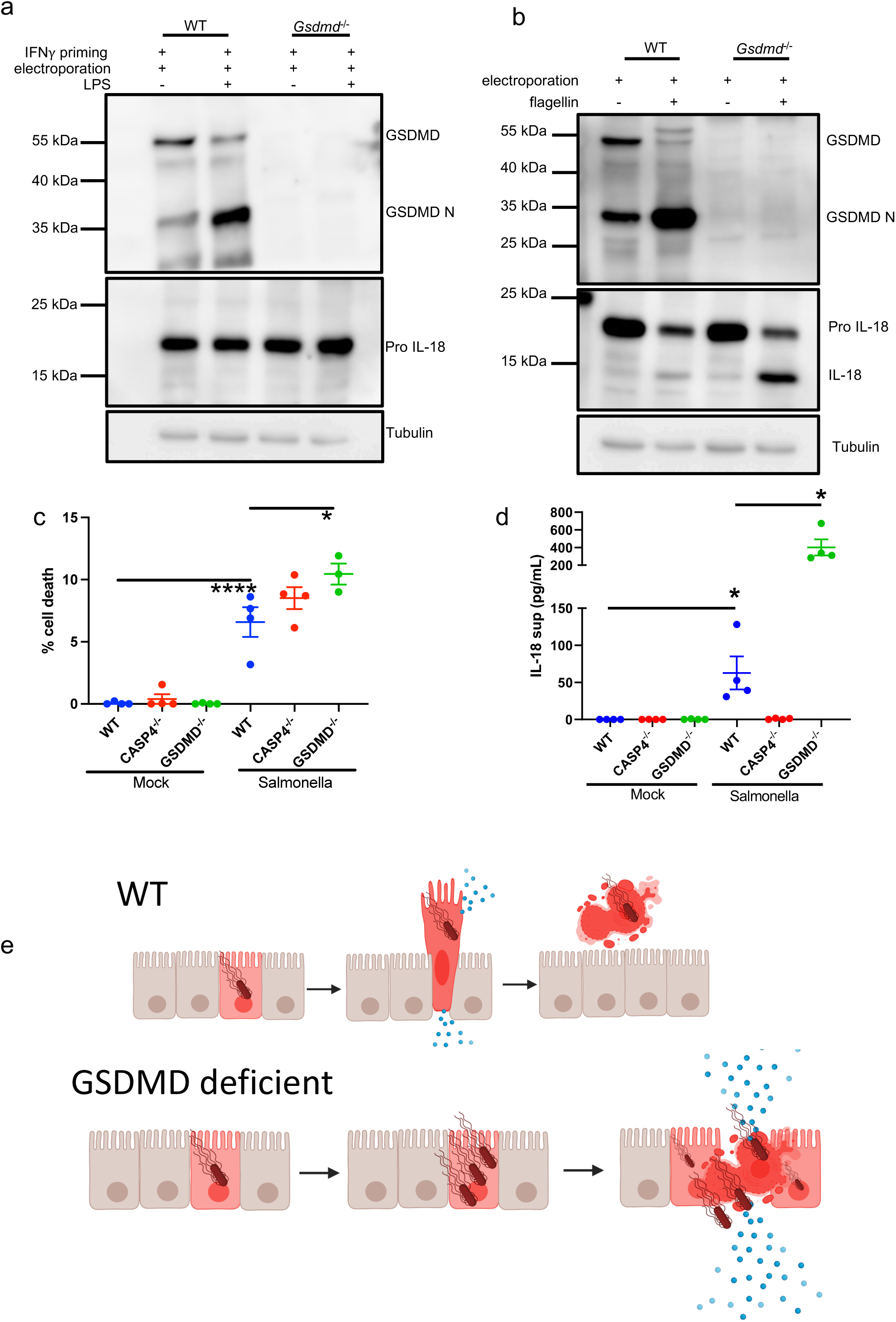
GSDMD regulates IL-18 production following murine NLRC4 activation and during *Salmonella* infection. Murine WT and *Gsdmd^-/-^* colonoids were electroporated with 5µg LPS (a) or 2.5µg of Flagellin (b). Cell lysates were collected 2 hours post electroporation and GSDMD cleavage and IL-18 production was measured by western blot (a, b). HCT116 cells were infection with WT Salmonella at an MOI of 50 for 16 hours. Cell death was measured by LDH cytotoxicity assay (c), IL-18 was measured by ELISA on supernatant (d). Graphical summary. (e). Representative of at least 3 experiments, data displayed as mean ± S.E.M. ****p<0.001, *p<0.05.

### Loss of GSDMD leads to IL-18 hypersecretion during *Salmonella* infection

We next sought to understand the physiological consequences of this feedback loop during infection. *Salmonella* induced inflammasome-independent cell death in infected HCT116 cells. Loss of caspase-4 inflammasome components led to increased cell death (Figure 8c), indicating that the caspase-4 inflammasome limits epithelial cell death during infection. *Salmonella* infection induced caspase-4 dependent IL-18 processing and secretion (Figure 8d). Importantly, loss of GSDMD increased IL-18 production and, due to increased cell death this was released into the supernatant (Figure 8d). Together, these data reveal that GSDMD pore formation regulates IL-18 production in both human and murine epithelial cells. In WT cells, GSDMD facilitates the controlled release of IL-18 and provides a signal to caspase-4 to terminate inflammasome signalling to control the production and release of cytokines. In the absence of GSDMD, IL-18 hyperproduction occurs. IL-18 remains trapped inside the cell, but upon the induction of GSDMD independent cell death pathways massive amounts of IL-18 are released (Figure 8e). In summary our data reveals a novel, epithelial-specific role for GSDMD pores in regulating inflammasome signalling and the production of IL-18.

## Discussion

Epithelial cell pyroptosis is a critical defense mechanism against invasive pathogens. In mice, NLRC4-dependent epithelial cell extrusion is critical for restricting *Salmonella*^7^, *Citrobacter rodentium*^25^ and *Shigella flexneri*^9^ invasion. This process is highly dependent on GSDMD^7,8^; however, little is known about what happens when GSDMD-mediated epithelial expulsion is inhibited or impaired. Inhibition of GSDMD is an immune evasion strategy engaged by multiple pathogens^26,27^, therefore there is a functional relevance to understanding how the epithelium responds to these conditions. In humans the caspase-4 inflammasome, rather than NLRC4 is the primary responder^10,11^ and far less is known about how caspase-4 coordinates pathogen restriction. Our work focused on understanding how the caspase-4 inflammasome functions in the intestinal epithelium. We reveal that in addition to controlling epithelial cell pyroptosis, GSDMD pore formation stimulates a negative feedback loop to inhibit caspase-4 activity and limit inflammatory cytokine production. In the context of infection, we propose that this results in the prompt removal of infected cells from the barrier and minimises induction of a wider immune response. However, when epithelial cell pyroptosis is inhibited, and infected cells cannot be ejected from the barrier, increased IL-18 heightens the inflammatory response. This is an effective way to modulate inflammation according to the degree of threat.

We observed that regulation of inflammasomes by GSDMD pores is largely epithelial-specific. Other studies have proposed mechanisms for how GSDMD regulates inflammasome activation in immune cells^28^. THP1 monocytes did not increase IL-18 production to the same degree as epithelial cells, with similar degrees of pore inhibition (Figure 2). We suspect that monocyte expression of NLRP3 accounts for this. NLRP3 activation amplifies GSDMD pore production leading to rapid pyroptosis and cellular lysis to trap the pathogen and recruit phagocytic machinery^6^. In this case cell death likely terminates inflammasome signalling. In epithelial barriers it seems the primary goal is to extrude infected cells and limit a wider immune response^7^. In this tissue, GSDMD activation can induce epithelial cell extrusion and simultaneously inhibit inflammasome activity. If the epithelium successfully extrudes the infected cell, the pathogen has been expelled and induction of a wider immune response is unnecessary. Conversely, if the epithelium is unable to extrude the cell, the pathogen represents a greater threat and induction of a wider immune response may be required to clear the pathogen. Interestingly, we observed that inhibition of NLRP3 did not lead to IL-18 hyperproduction in immune cells. Indeed, inhibition of NLRP3 inhibited IL-18 production (Supplementary Figure 1). This likely indicates that in immune cells IL-18 production is dependent on caspase-1. The reason for differences in IL-18 processing capacity between cell types requires further investigation.

We additionally found that full length caspase-4 processes IL-18. Within the field, the consensus is that caspase auto-processing is required for catalytic activity against substrates^19,20,29^. However, auto-processing requires induction of catalytic activity, and oligomerization serves as the signal for induction of auto-processing capacity. Thus, it is plausible that oligomerization is sufficient for generation of catalytic activity. A similar theme has been seen in other caspase biology. Non-cleavable caspase-1 has activity against GSDMD^30^. Similarly, full length caspase-8, in a heterodimeric complex with FLIP_L_ or in a homodimeric complex with itself is sufficient to generate catalytic activity^31^. Moreover, LPS-induced oligomerization of caspase-4 is sufficient to generate activity against the artificial caspase substrate Z-VAD-AMC^32^. Studies reporting caspase-4 cleavage have examined caspase activation in WT cells^14,20,33^ and in overexpression systems^34^. We similarly see caspase-4 cleavage under these conditions, and in all cases caspase-4 processing was associated with reduced IL-18 production. We therefore propose that caspase-4 cleavage terminates or limits IL-18 processing, and that the caspase-4 cleavage fragment is a by-product of inflammasome activation, rather than the active species. Trapping cleaved caspase-4 inside cells had no effect on IL-18 or GSDMD processing (Figure 6e), further ratifying this idea. It is also interesting to note that the p31 fragment observed in THP1 cells (Figure 3c, d) and in our HEK overexpression system (Figure 5f) was not due to caspase-4 autocatalytic activity. Inhibition of NLRP3 prevented p31 generation in THP1 cells and the p31 fragment was observed in HEK cells expressing the catalytically dead caspase-4 mutant C258A. This indicates that other enzymes may cleave caspase-4 and hence alter caspase-4 activity. Interestingly, the cleavage profile observed in HCT116 cells was different to that observed in THP1 cells (Figure 3a, c). This may indicate different mechanisms of cleavage in a cell-specific manner which may be engaged in a pore dependent manner to regulate caspase activity.

While we observed that GSDMD pore formation was required for caspase-4 cleavage, we have not determined how the pore relays this signal. Calcium influx controls activation of the ESCRT pathway^15^, rearrangement of F-actin^35^ and the openness of GSDMD pores^36^. Therefore, calcium is an attractive target. How calcium would induce caspase-4 cleavage is unknown. Data is emerging showing plasma membrane pore independent functions of GSDMD. For example, GSDMD permeabilizes mitochondria and lysosomes^37^, and studies are reporting GSDMD N-terminus can relocate to the nucleus^38^. We cannot discount that a pore-independent mechanism involving the GSDMD N-terminus may regulate caspase-4 activity. More knowledge is needed on how to control GSDMD N-terminal localisation to accurately determine if pore-independent mechanisms control inflammasome activation.

In summary we have identified a novel mechanism for GSDMD-dependent regulation of the production of IL-18 in the intestinal epithelium. This mechanism appears to be specific to epithelial cells but occurs across inflammasomes and species. Additionally, we provide the first evidence to our knowledge that caspase-4 directly processes IL-18. We moreover demonstrate that full-length caspase-4 has the greatest IL-18 processing capacity. Our observation that GSDMD regulates inflammasome cytokine production alludes to a novel pathway through which inflammasome signalling can be modulated.

## Author Contributions

S.E.G, J.K.B, D.J.P and S.K designed the experiments. L.L and Y.T performed THP1 experiments, N.W performed the flow cytometry assays and analysis, C.K created CRISPR knockout cells. J.K.B performed all other experiments and analysis.

---

**Supplementary Figure 1:**
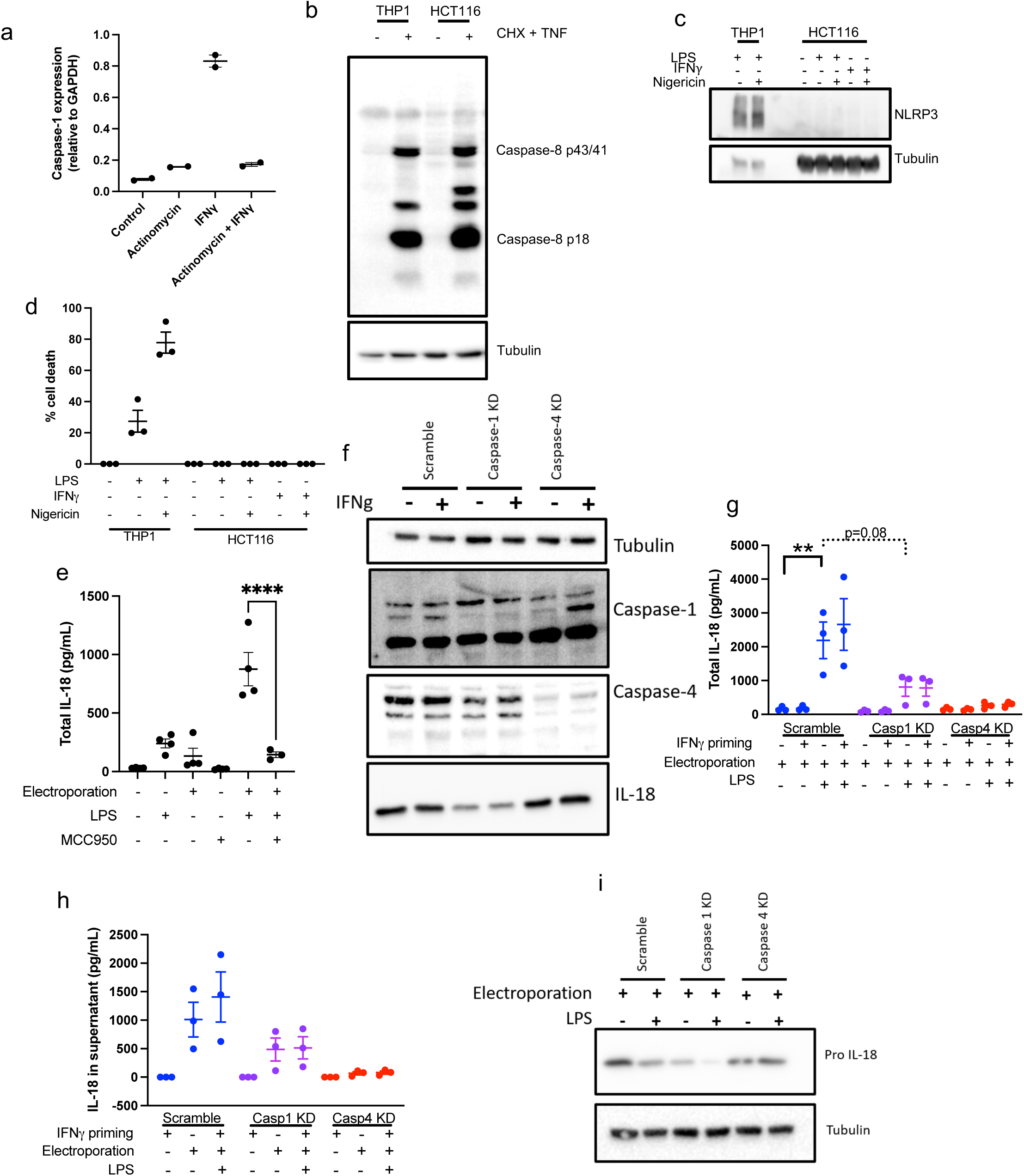
(a) HCT116 cells were treated with Actinomycin D then primed with IFNγ. (b) THP1 or HCT116 cells were treated with cycloheximde at concentration in Figure 1h and costimulated with TNF to show CHX concentration working. (c-d) THP1 or HCT116 cells were treated with 100ng/mL LPS or 10ng/mL IFNγ then simulated with 5ug/mL nigericin for 30 minutes. NLRP3 levels were assessed by western blot (c), cell death was measured by LDH cytotoxicity assay (d). (e) THP1 cells were stimulated with LPS as in Figure 2e, cells were lysed into supernatant and IL-18 levels were measured by ELISA. (f-i) Indicated caspases were silenced by shRNA in C2bbe1 epithelial cells. Cells were primed with 10ng/mL IFNγ to prime caspase-1 (f,-h). Cells were electroporated with 2µg LPS and IL-18 was measured in cell free supernatants (g), or cells were lysed into supernatants and total IL-18 was measured by ELISA (h). Pro IL-18 in whole cell lysates was measured by western blot (i). Representative of at least 3 experiments, data displayed as mean ± S.E.M. **p<0.01, ****p<0.001.

